# EEG Dynamical Features during Variable-Intensity Cycling Exercise in Parkinson’s Disease

**DOI:** 10.1101/2025.01.15.633102

**Authors:** Zahra Alizadeh, Emad Arasteh, Maryam S. Mirian, Matthew A. Sacheli, Danielle Murray, Silke Appel-Cresswell, Martin J. McKeown

## Abstract

1.

**Background:** Exercise is increasingly recognized as a beneficial intervention for Parkinson’s disease (PD), yet the optimal type and intensity of exercise remain unclear. This study investigated the relationship between exercise intensity and neural responses in PD patients, using electroencephalography (EEG) to explore potential neural markers that could be ultimately used to guide exercise intensity.

**Method:** EEG data were collected from 14 PD patients (5 females) and 8 healthy controls (HC) performing stationary pedaling exercises at 60 RPM with resistance adjusted to target heart rates of 30%, 40%, 50%, 60%, and 70% of maximum heart rate. Subjects pedaled for 3 minutes at each intensity level in a counterbalanced order. Canonical Time-series Characteristics (Catch-22) features and Multi-set Canonical Correlation Analysis (MCCA) were utilized to identify common profiles of EEG features at increasing exercise intensity across subjects.

**Results:** We identified a statistically significant MCCA component demonstrating a monotonic relationship with pedaling intensity. The dominant feature in this component was Periodicity Wang (PW), related to the autocorrelation of the EEG. Analysis revealed a consistent trend across features: six features increased with intensity, indicating heightened rhythmic engagement and sustained neural activation, while three features decreased, suggesting reduced variability and enhanced predictability in neural responses. Notably, PD patients exhibited more rigid, consistent response patterns compared to healthy controls (HC), who showed greater flexibility and variability in their neural adaptation across intensities.

**Conclusion:** This study highlights the feasibility of using EEG-derived features to track exercise intensity in PD patients, identifying specific neural markers correlating with varying intensity levels. PD subjects demonstrate less inter-subject variability in motor responses to increasing intensity. Our results suggest that EEG biomarkers can be used to assess differing brain involvement with the same exercise of increasing intensity, potentially useful for guiding targeted therapeutic strategies and maximizing the neurological benefits of exercise in PD.

## 2. Introduction

PD is a neurodegenerative disease associated with the accumulation of alpha-synuclein and loss of dopamine-producing neurons, particularly in the substantia nigra (Calabresi et al., 2023). Symptoms of this dopaminergic neuron loss include bradykinesia, muscle rigidity, and tremors, in addition to a host of nonmotor symptoms (Licen et al., 2022). Current treatments such as medication and deep brain stimulation (DBS) can alleviate its symptoms but do not fundamentally alter the course of the disease and vary in effectiveness depending on the individual (Ridgel et al., 2015). Consequently, it is crucial to explore new therapeutic approaches which can have rehabilitation potential without side effects.

Numerous studies have investigated the potential of exercise in improving symptoms of various neurological problems including neurodegenerative diseases such as Parkinson’s disease (Langeskov-Christensen et al., 2024), and Alzheimer’s (Valenzuela et al., 2020). (Valenzuela et al., 2020) shed light on the advantageous role of exercise in mitigating Alzheimer’s disease hallmarks. Exercise has also had a positive impact on balance in patients with stroke, PD, and multiple sclerosis, (for a systematic review see: (Salari et al., 2022)). Exercise has also been beneficial in psychiatric diseases such as anxiety (Pontifex et al., 2021), and Obsessive Compulsive Disorder (OCD) (Abrantes et al., 2019). In a metaanalysis, (Ensari et al., 2015) showed that exercise provides immediate relief from state anxiety, and (Pontifex et al., 2021) demonstrated its ability to alleviate cognitive impairments associated with anxiety.

Cycling appears to be a particularly beneficial exercise for individuals with Parkinson’s disease (PD). (Alberts et al., n.d.) noted improvements in a person living with Parkinson’s disease (PwP) within two days after a tandem cycling expedition. Follow-up studies (Ridgel et al., 2012) (Ridgel et al., 2009) (Ridgel et al., 2015) revealed significant benefits, including a 38% reduction in tremors and a 28% improvement in bradykinesia after eight weeks of forced tandem cycling. Dynamic high-cadence cycling also showed motor improvements, while other research (Snijders et al., 2012)(Licen et al., 2022) highlighted the persistence of cycling ability in more advanced disease, even in those with severe freezing of gait. The repetitive nature of cycling poses a lower physical challenge and may help stabilize abnormal neural oscillations, making it a promising rehabilitative option for PD patients (Licen et al., 2022).

Despite the well-documented beneficial effects of exercise on the brain function, gaps remain in understanding how factors like duration, type, and intensity influence these outcomes. Most studies have focused on comparing pre- and post-exercise brain states (Moraes et al., 2007) (Gutmann et al., 2015). However, monitoring EEG *during* exercise could provide valuable insights into how these variables interact (Enders et al., 2016) (Bailey et al., n.d.), potentially enhancing our ability to optimize exercise-based therapeutic interventions.

Recent research has focused on leveraging dynamic features from brain recordings like EEG to better understand neurodegenerative diseases (Cacciotti et al., 2024). These features offer critical insights into the temporal fluctuations of brain connectivity, which may assist in interpreting symptom progression, and mechanisms behind cognitive and motor impairments. (Lubba et al., 2019) proposed 22 key temporal features, known as Canonical Time-series Characteristics (Catch-22), from an initial set of 4,791. These features effectively capture diverse aspects of temporal dynamics of various time series, including EEG, that capture extreme events, autocorrelation, symbolic patterns, and scaling behaviors, while reducing redundancy, making them broadly applicable across time-series analyses.

In this study, we applied Catch-22 dynamic EEG features to assess the effects of pedaling intensity on brain activity in individuals with Parkinson’s disease (PD) and healthy controls. Participants pedaled at five different intensity levels while their EEG data were collected. Using Multi-set Canonical Correlation Analysis (MCCA) (Fu et al., 2019), we identified linear combinations of EEG channels that exhibited consistent trends across participants, highlighting key neural responses as pedaling intensity increased.

## 3. Methods

### Experimental Protocol

In this study, 14 PD and 8 healthy control (HC) subjects were recruited from the Movement Disorders Clinic at the Pacific Parkinson’s Research Centre, Vancouver, Canada. All subjects provided written, informed consent, and all research was reviewed and approved by the appropriate Ethics Boards. To optimize exercise tolerance, subjects took their usual anti-parkinsonian medication. People with stroke, severe cardiovascular disease, and other significant neurological or psychiatric disorders were excluded from the study. More information about PD subjects is in Table 1.

**Table 1.** Demographic information about the subjects included in this study.

| Variables | Values |
| --- | --- |
| Age | $60.66 \pm 7.81$ |
| Male/Female | 9/5 |
| Hoehn-Yahr | 1-3 |
| Regular exercise | >100 minutes per week over 3 months |
| Montreal Cognitive Assessment Score | >24<br>($27.53 \pm 1.92$) |
| Beck’s Depression Score | <13<br>($2.27 \pm 3.63$) |
| Beck’s Anxiety Score | <9<br>($3.67 \pm 3.2$) |
| Medication | Beta-blocker “Off”<br>Anti-Parkinson “On” |
| Handedness (Right/Left) | 11/4 |

Participants pedaled on a stationary bicycle, with the seat height individually adjusted for comfort. Each session began with a 5-minute warm-up to ensure safety. Subjects pedaled at intensities of 30%, 40%, 50%, 60%, and 70% of their maximum heart rate (calculated as 0.85 * [220 - age]) while maintaining a steady pace of 60 RPM. Resistance was adjusted to vary intensity, with 3-minute intervals followed by 3-minute recovery periods. Intensity intervals were pseudo-randomly assigned, except for 70%, which never came first. The 30- minute intervention concluded with a 5-minute cool-down.

EEG was recorded using nine electrodes at standard locations F3, Fz, F4, C3, Cz, C4, P3, Pz, and P4, targeting frontal, central, and parietal regions critical for monitoring brain activity during exercise. A stretchable head cap with Ag-AgCl electrodes (ANT Wave Guard cap, Advanced Neuro Technology B.V.) was used, with the reference electrode placed near the vertex between Cz and CPz. The ground electrode was positioned between Fz and FPz on the frontal scalp, and electrode impedances were maintained below 10 kΩ to ensure signal quality.

### 3.1. Preprocessing

We performed a number of careful steps to remove the artifact from the EEG. First, the common average reference (CAR) technique was applied, followed by high pass FIR filtering, low pass FIR filtering, and notch filtering to remove frequencies below 0.1, above 100, and 60 Hz line noise. A recursive least squares (RLS) algorithm was applied to remove eye-related (ocular) artifacts, and an adaptive filter was used in this method to remove eye blinks from the EEG signal by adjusting its coefficients in each iteration according to the difference between the EEG signal and the EOG signal. The automatic artifact rejection (AAR) plugin was used in EEGLAB (MATLAB 2018b) to reject this artifact. Then, the EMG artifact was removed with principal component analysis (PCA). In the last step, another remaining artifact source was removed with the help of the IClabel plugin of EEGLAB, with two of the nine components removed.

### 3.2. Feature Extraction Using Catch-22

We utilized the Catch-22 package, which allows for the extraction of 22 distinct features from each EEG channel, as listed in Table S1. The duration for feature extraction was specifically set to one second, corresponding to one complete cycle of pedaling (60 RPM). By synchronizing the feature extraction window with the pedaling cycle, we effectively isolated and minimized the impact of pedaling frequency on the EEG signals, ensuring that the extracted features more accurately reflected the underlying brain activity rather than the mechanical motion.

### 3.3. MCCA-Based Extraction of Intensity-Linked Neural Signatures

MCCA extends traditional CCA by analyzing relationships across multiple datasets. It identifies optimal linear combinations of variables (canonical variates) that maximize correlations between datasets. In this study, MCCA was applied to EEG data from 14 participants in a two-step process (Fig. 1).

**Fig. 1.**
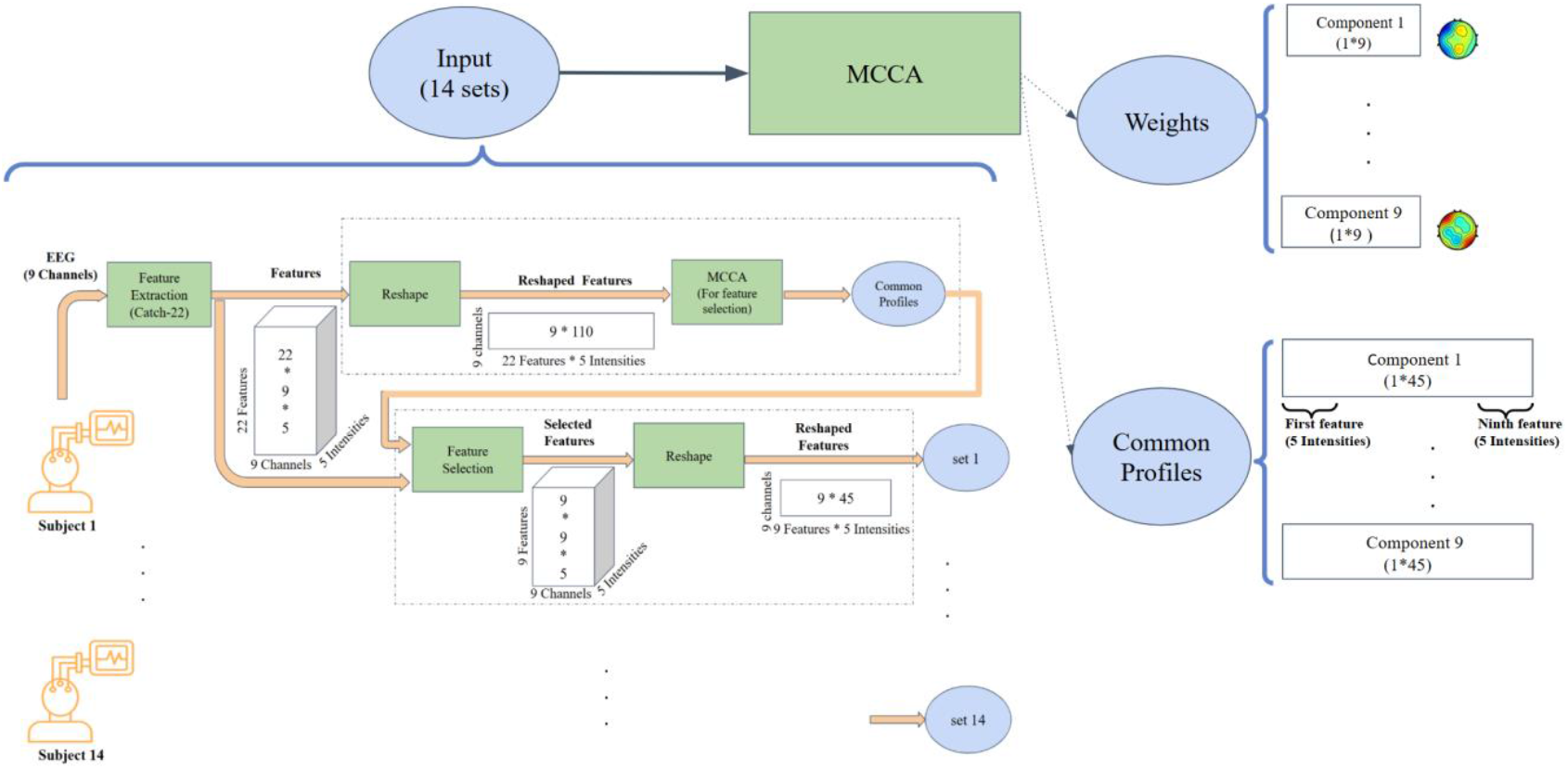
The pipeline displays the identification process of the most consistent features across EEG channels, highlighting those with the strongest relationship to exercise intensity.

#### 3.3.1. First Round: Feature Selection

In the first round (upper dashed rectangle in Fig. 1), the data matrix for each subject was structured as 9 EEG channels by 110 observations (22 features across five intensity levels). This round served as a feature selection step, aiming to identify features that consistently represented shared neural dynamics across EEG channels of all subjects. Nine canonical components were extracted (corresponding to the number of EEG channels), and features with the highest consistency across subjects were identified for further analysis.

#### 3.3.2. Second Round: Detecting Neural Signatures

Building on the results above, the second round of MCCA focused on detecting the neural signatures of exercise intensity. Informative features selected from the first round, evaluated for their consistency across subjects within components 2–5 (as detailed in the results section), were used to refine the data matrices. These matrices, structured as 9 channels by 45 observations (9 features across 5 intensities), enabled more targeted analysis. This two-step approach ensured improved interpretability and computational efficiency by concentrating on features most relevant to exercise-induced neural dynamics.

### 3.4. Weighted Signal Analysis and Wavelet Transformation

To integrate the multichannel EEG data, a weighted aggregate signal was computed using channel weights derived from the MCCA analysis. To capture critical events in both the time and frequency domains, the Wavelet Transform was employed. Before transformation, EEG signals were averaged over one-second windows (corresponding to a single pedaling cycle at 60 RPM) to minimize noise and artifacts. This averaging step ensured that the signals reflected underlying neural activity rather than transient noise or mechanical motion.

## 4. Results

### Artifact Identification via Principal Component Analysis

Consistent with prior studies (De Cheveigné et al., 2019) the first component of MCCA may still relate to residual artifacts (Jiang et al., 2019) as illustrated in Fig. 2 for one subject. We therefore excluded the first component and concentrated our further investigations on components 2-4.

**Figure. 2.**
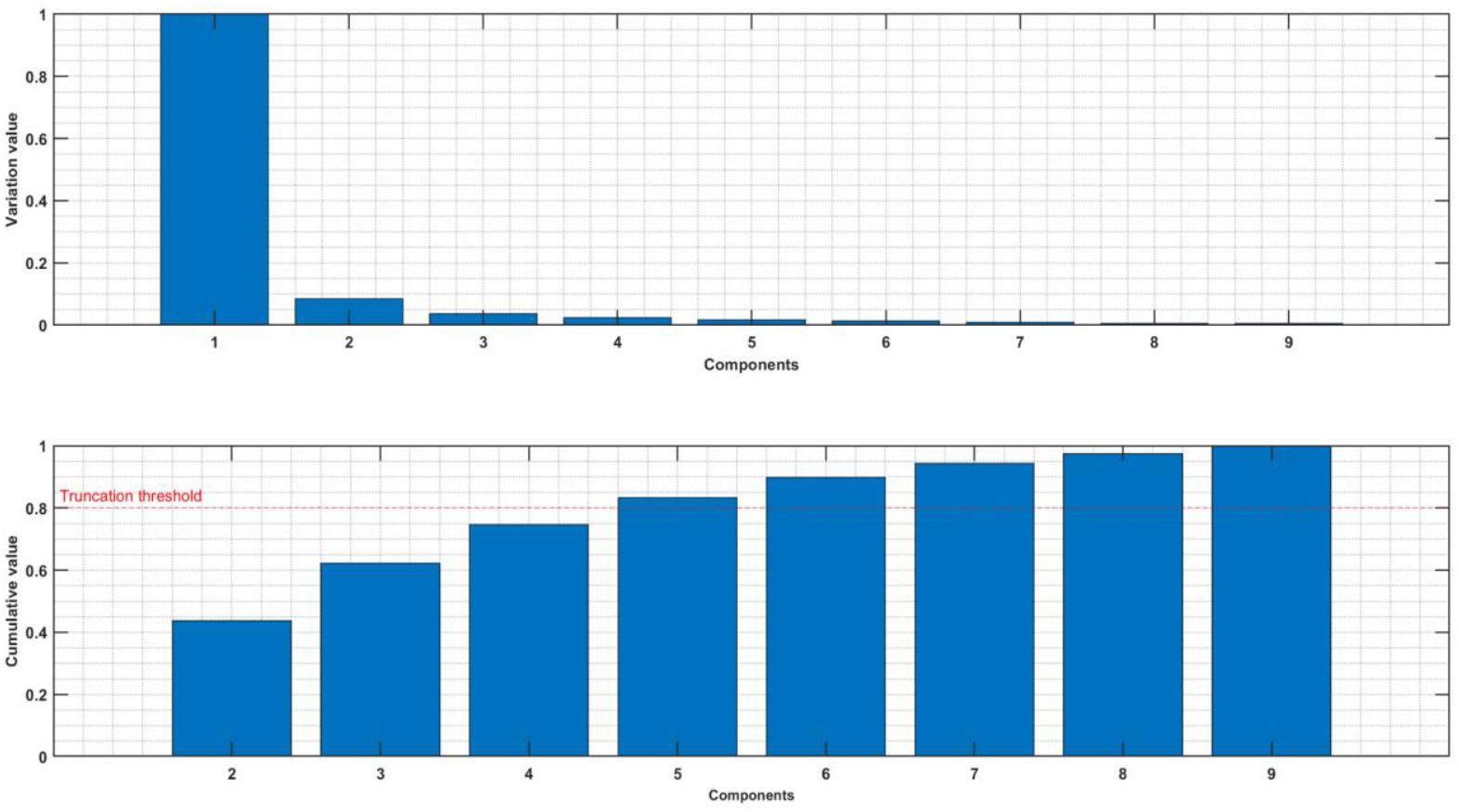
Upper panel: Normalized variance of each component of MCCA. Lower panel: Normalized cumulative variance of components 2-9.

### 4.1. MCCA-Based Extraction of Intensity-Linked Neural Signatures

MCCA analysis at the first round demonstrated nine features that exhibited high consistency across subjects within the meaningful components 2, 3, and 4. The feature values were normalized separately to obtain a better visual representation. Nine out of the 22 features exhibited consistent behavior across subjects. A detailed description of these nine consistent features is presented in Table 2. The results for component 3 are shown in Fig. 3 and Fig. 4, with supplementary Fig. S1 showing the control group for comparison. Six features, as mentioned in the second column of Table 2, showed an increasing trend with rising exercise intensity, while three features showed a decreasing trend.

**Table 2:**
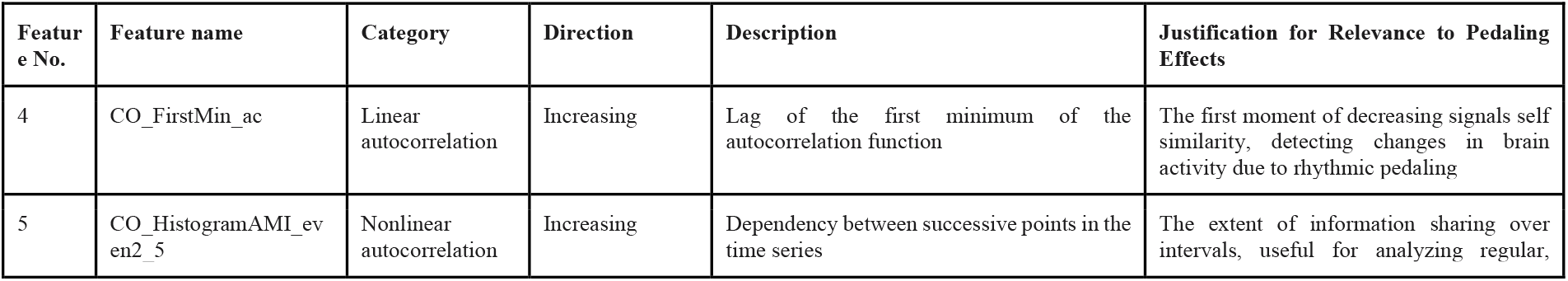
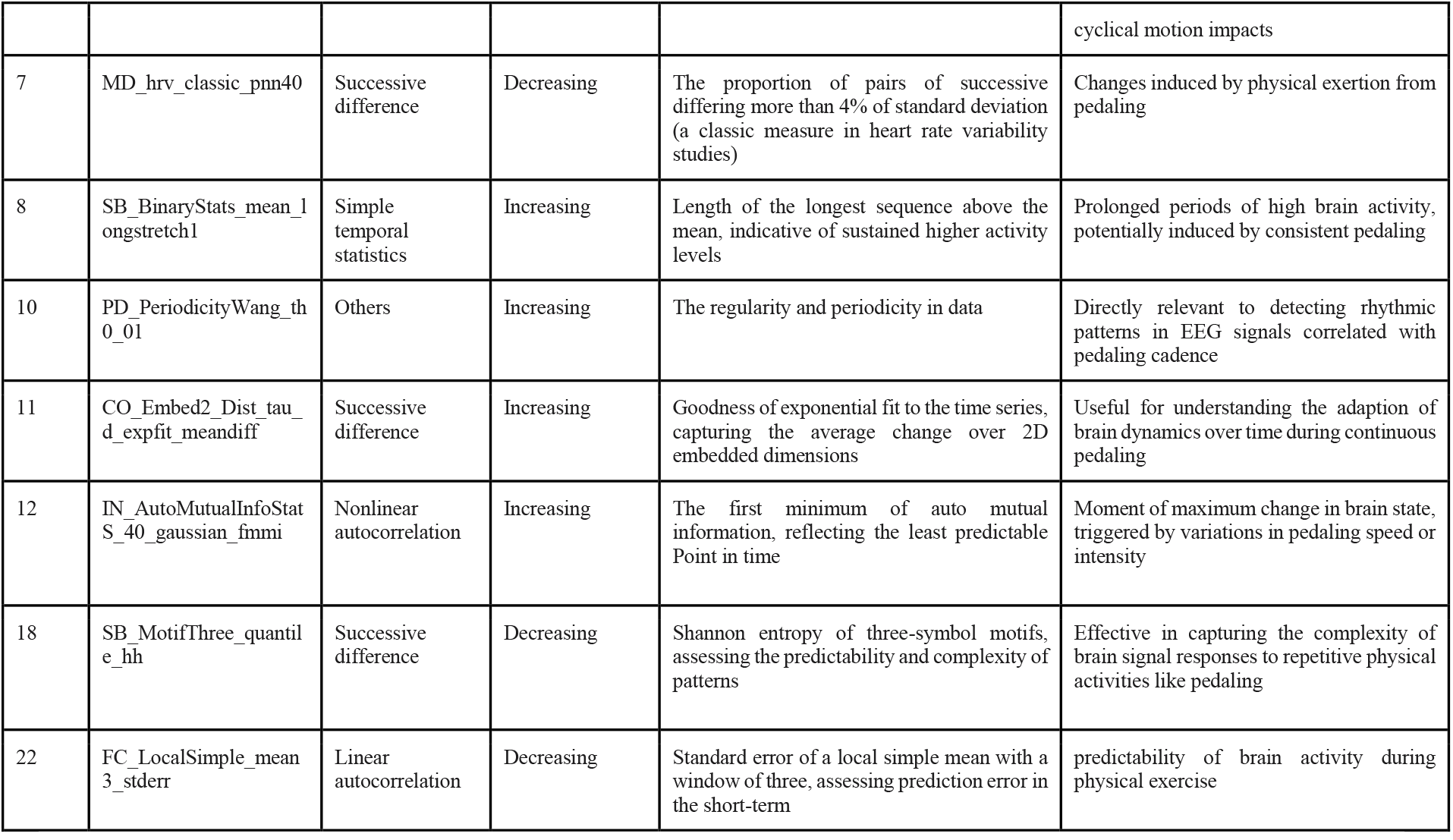
Most consistent features of the Catch-22 package. Feature numbers and names are mentioned based on numbers listed in (Lubba et al., 2019).

**Figure. 3.**
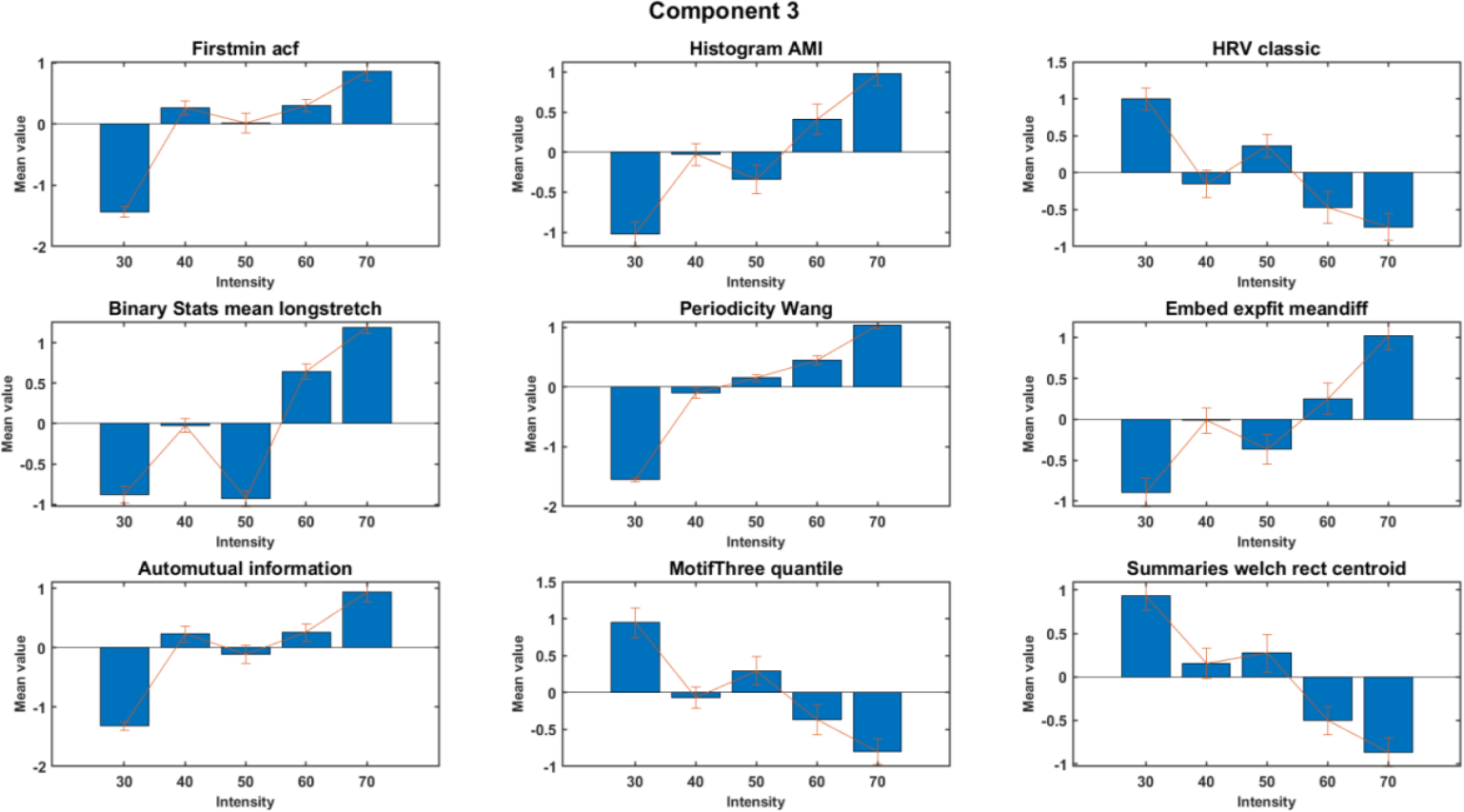
Trend across intensity for features of component 3 of PD subjects. The figure shows the aggregated (normalized) mean and standard deviation across subjects.

**Figure. 4.**
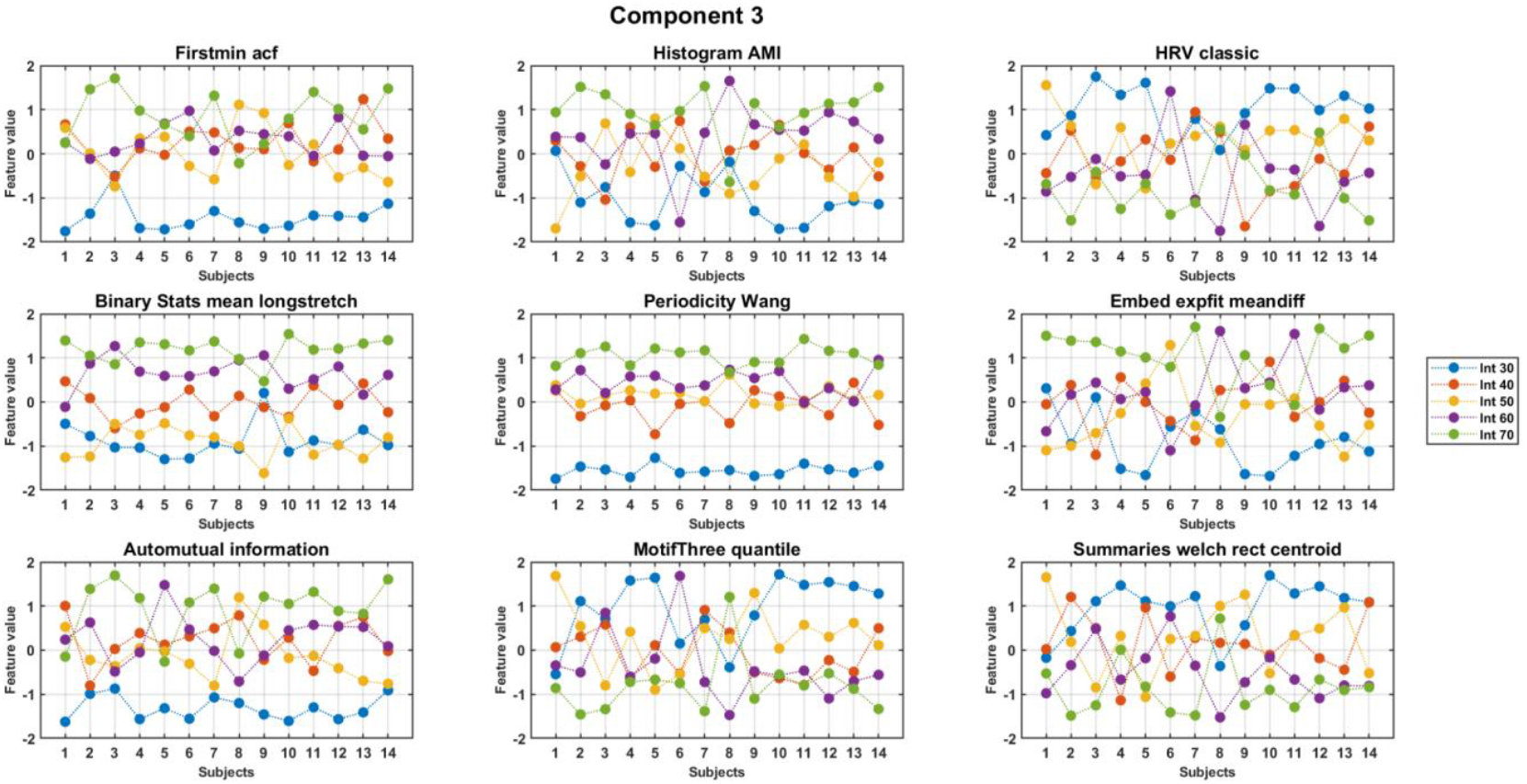
Trends across intensity for features of component 3 of PD subjects. The figure displays (normalized) values of features for individual subjects (x-axis) for each feature.

Moreover, by focusing on the weighted vector of EEG data computing by multiplying EEG by the weights of MCCA, we observed a distinct cluster of harmonics that was concentrated around the alpha band, as depicted in the supplementary section in Fig. S4.

### 4.2. Post-hoc Analysis: investigation of the Difference between PD and HC

To evaluate differences in pedaling intensity and compare individuals with Parkinson’s disease (PD) to healthy controls (HC), we conducted a repeated measures Analysis of Variance (ANOVA). We examined both the within-subject factor of pedaling intensity and the between-subject factor of group classification (PD vs. HC) to understand how features vary under different conditions. This analysis considered pedaling intensity as a within-subject factor and group classification (PD vs. HC) as a between-subject factor. Results showed significant intensity interactions within the PD group (10.25 < F < 514.5, 0 < p < 0.004) and significant differences between groups (8.79 < F < 533.77, 0 < p < 0.007). HC subjects exhibited greater variability across intensities, while PD participants showed more consistent, predictable feature changes. Details are provided in Supplementary Table S2.

## 5. Discussion

This study offers new insights into the link between exercise intensity and brain dynamics in Parkinson’s disease (PD), highlighting distinct neural activity patterns during pedaling. The key finding is the identification of consistent EEG features, derived through Multi-set Canonical Correlation Analysis (MCCA), that reliably track changes in pedaling intensity. These measurable and robust neural responses across subjects present promising potential biomarkers for tailoring and optimizing exercise-based interventions for PD management.

If increasing exercise intensity can alter brain activity, it may further activate existing neural regions or recruit new ones, both of which have been described as compensatory mechanisms in Parkinson’s disease (PD) [ref]. By applying the Multi-set Canonical Correlation Analysis (MCCA) to identify consistent Catch-22 EEG features across subjects, we allowed individual channel combinations to vary from subject-to-subject. Six features in Component 3 increased with intensity (FirstMin_acf, HistogramAMI_even_25, BinaryStats_meanlongstretch, PeriodicityWang, Embed2_Dist_expfit_meandiffand and AutoMutualInfoStats), reflecting enhanced rhythmic coordination, prolonged engagement, and adaptive responses. Conversely, three features (Hrv_classic, MotifThree_quantile, and LocalSimple_mean3stderr) decreased, indicating reduced variability and increased neural efficiency, aligning with the demands of higher-intensity exercise. These results align with prior work indicating that increased exercise intensity influences neural oscillations, in the alpha frequency range (Ciria et al., 2018). Prior work has shown that cycling has been shown to both suppress beta-band local field potentials from Deep Brain Stimulation electrodes but also enhance narrowband power increases around 18Hz (Storzer et al., 2017).

We observed reduced variability and flexibility in neural responses to increasing exercise intensity in individuals with Parkinson’s disease (PD). In PD, beta-band oscillations are excessively synchronized, with more consistent waveform features, such as sharper peaks and greater steepness asymmetry, compared to healthy controls (Jackson et al., 2019). In PD patients with Freezing of Gait, higher EEG amplitude synchronization across frequency bands (theta, alpha, beta, and gamma) has been observed between different brain regions, regardless of the motor task (Asher et al., 2021). These observations suggest that subjects with PD have impaired metastability, a critical property of the brain’s dynamics. Metastability reflects the brain’s ability to balance stability and flexibility, supporting efficient information processing, cognitive function, and adaptability. In the context of exercise, it enables the brain to transition between varying intensity levels, expanding its dynamic repertoire (Bapat et al., 2024) and supporting the functional states needed for diverse motor and cognitive activities.

Our analysis primarily relied on linear methods, such as Multi-set Canonical Correlation Analysis (MCCA), which may only partially capture the complexity of neural responses during exercise. Future studies should explore nonlinear approaches to better detect exercise-induced neural changes and uncover subtler patterns of brain activity. It is worth noting, however, that the Catch-22 package already includes several nonlinear measures, which offer a valuable starting point for these investigations. Additionally, the use of only nine EEG electrodes limited the spatial resolution of the recorded brain activity, suggesting that future research could benefit from higher-density EEG arrays or multimodal neuroimaging techniques, such as combining EEG with fMRI. Further research could also assess the consistency of the identified features across varying pedaling intensities in a larger cohort, ensuring their robustness and relevance to motor performance and clinical outcomes. The potential of the Periodicity Wang feature as a biomarker could be explored using other modalities, such as fMRI and ECG, to determine its broader applicability across neural and physiological contexts.

## 6. Conclusion

This study highlights the potential of EEG-derived features to monitor and assess exercise intensity in Parkinson’s disease (PD) patients. Among these, the Periodicity Wang (PW) feature, which measures the autocorrelation of neural dynamics, showed a strong association with changes in pedaling intensity. By applying Catch-22 features and Multi-set Canonical Correlation Analysis (MCCA), we observed consistent neural responses across subjects at different intensity levels. Some features exhibited increasing trends, indicating enhanced rhythmic coordination and sustained neural activation, while others displayed decreasing trends, reflecting reduced variability and a transition to more stable, predictable neural patterns. These findings offer valuable insights for optimizing exercise-based interventions in PD and highlight the potential for identifying neural biomarkers to guide future therapeutic strategies.

## 7. Acknowledgment

This work was supported by resources made available through the Dynamic Brain Circuits cluster and the NeuroImaging and NeuroComputation Centre at the UBC Djavad Mowafaghian Centre for Brain Health (RRID SCR_019086). MJM was supported by the John Nichol Chair in Parkinson’s Research. SAC is supported by the Marg Meikle Professorship for Parkinson’s Disease Research through the Pacific Parkinson’s Research Institute.

## 8. Code/Data Availability

The GitHub repository, including the codes of this paper, can be accessed here: https://github.com/zahraalz3523.

All data produced in the present study are available upon request to the authors.

## 11. Supplementary

Supplementary 1: A description of all Catch-22 features is brought in Table S1.

**Table S1:**
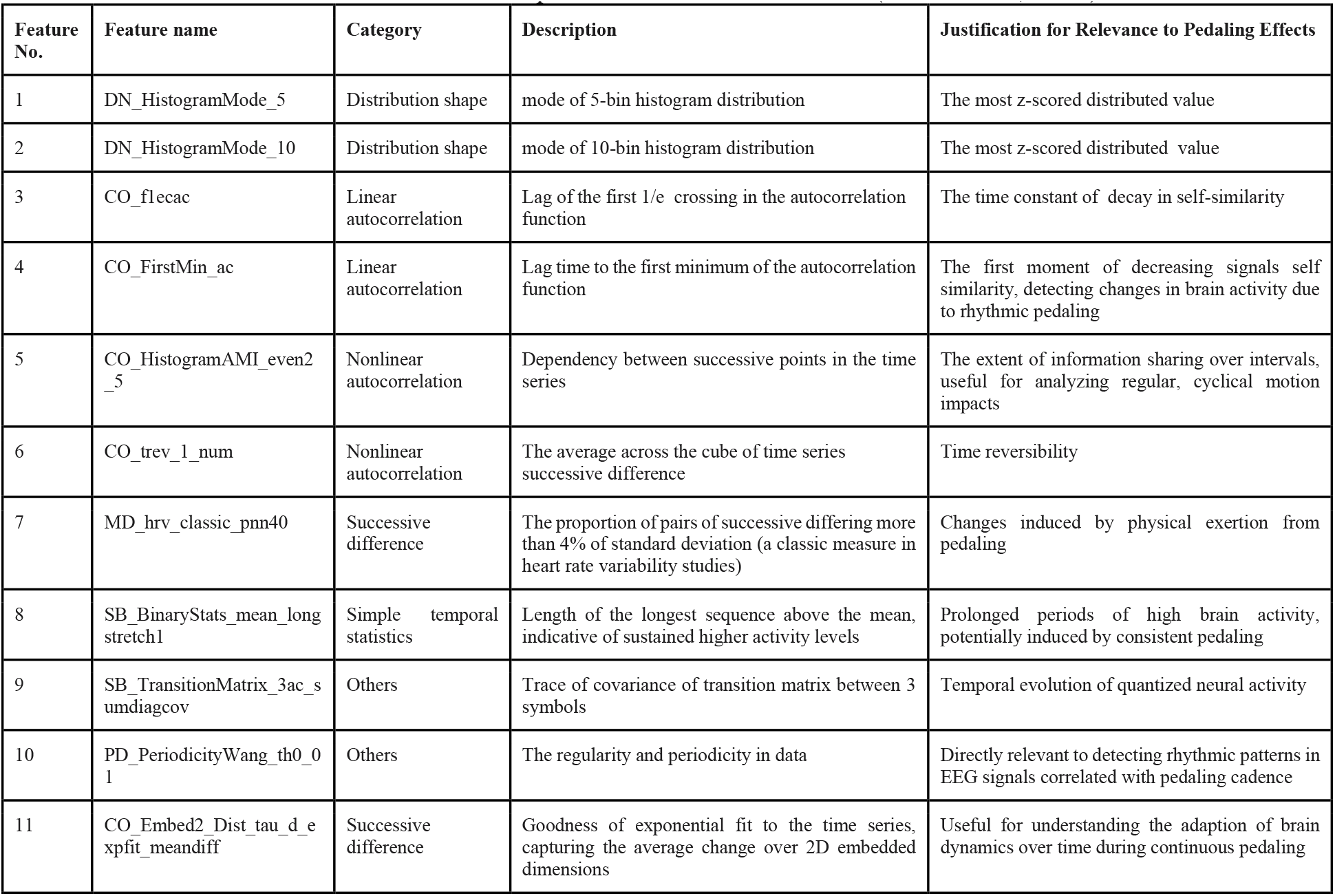
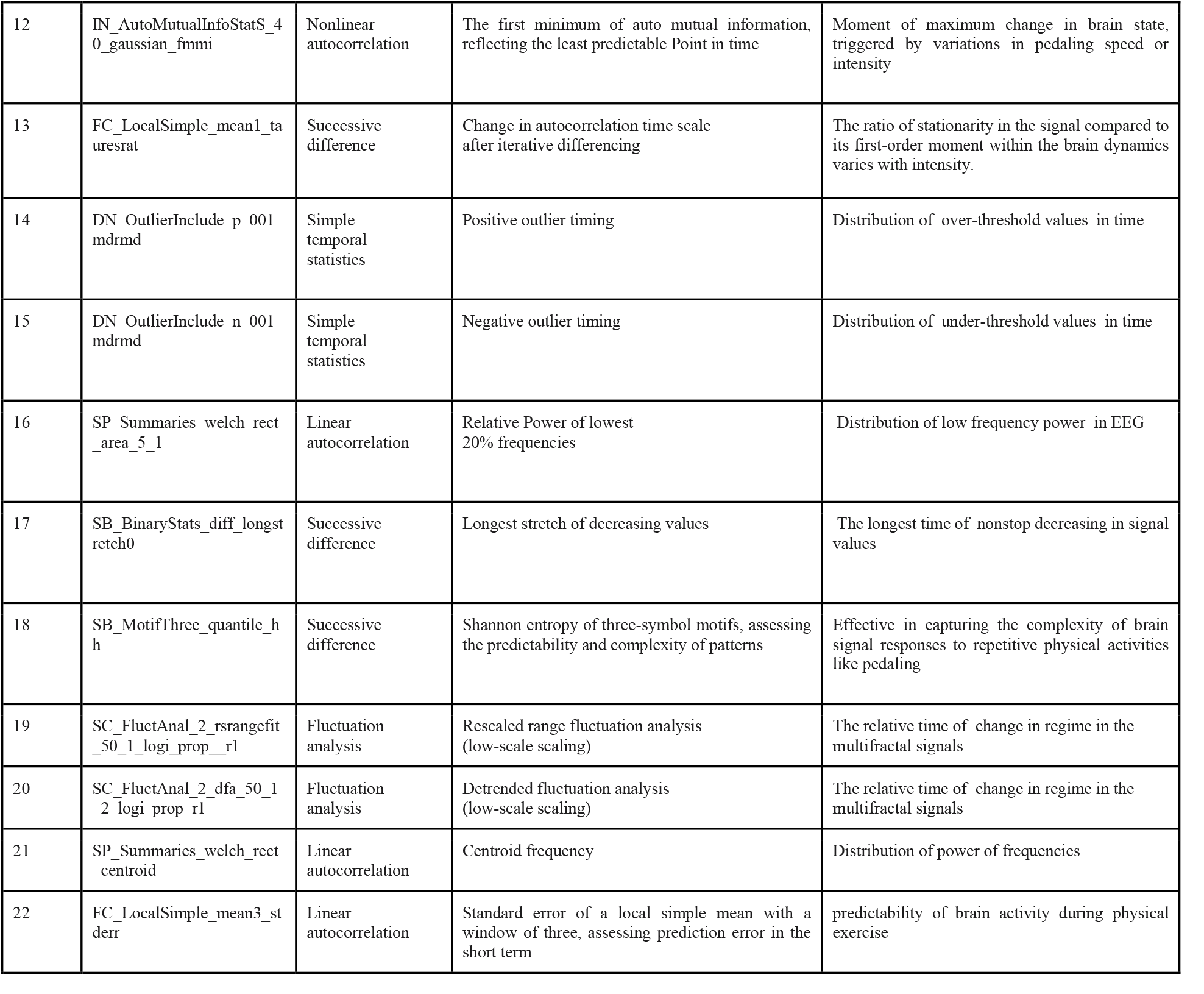
Features and interpretation of Catch-22 features (Lubba et al., 2019)

Supplementary 2: We also applied MCCA to the control group to establish a baseline representation of the common patterns within a healthy population. This analysis enables a direct comparison with the PD group, allowing us to identify distinct differences and shared features between the two groups. By doing so, we can better understand the variations introduced by pathological conditions and ensure that our findings are robust and grounded in meaningful contrasts. The results of this analysis for the control group are brought in Fig. S1.

**Fig. S1.**
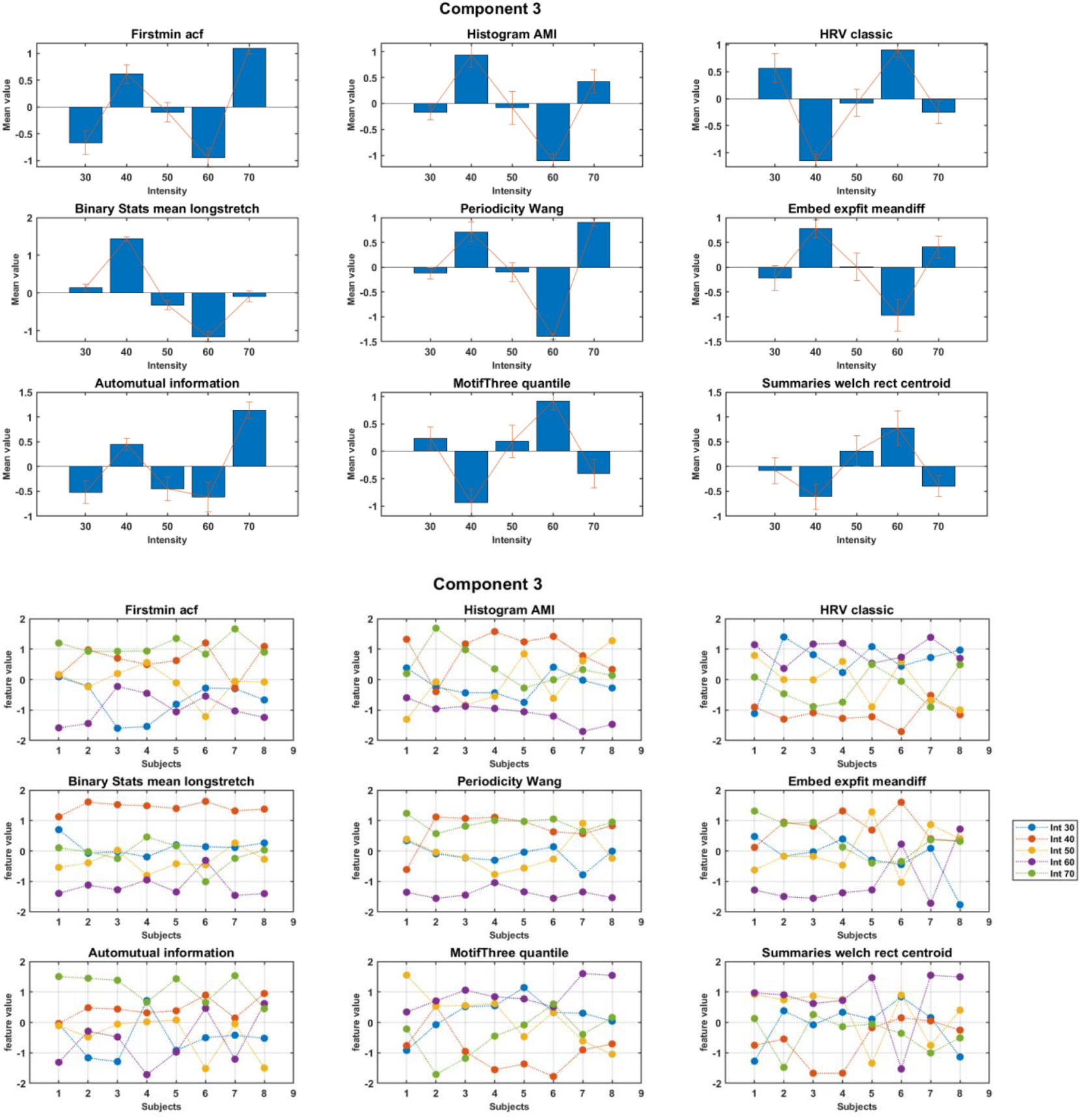
Trend across intensity for features in component 3 for control group. The upper figure shows the aggregated (normalized) mean and standard deviation across subjects. The lower figure shows feature values for all subjects

Supplementary 3: We conducted two rounds of MCCA. In the first round, including 22 features, we analyzed Components 2–4 to identify most consistent features over subjects. Based on this analysis, we selected 9 features for the second MCCA. These features were then used to perform the second round of MCCA. Results of component 3 were brought in the main transcript, and components 2 and 4 are brought in fig S.2 and S.3.

**Fig. S2.**
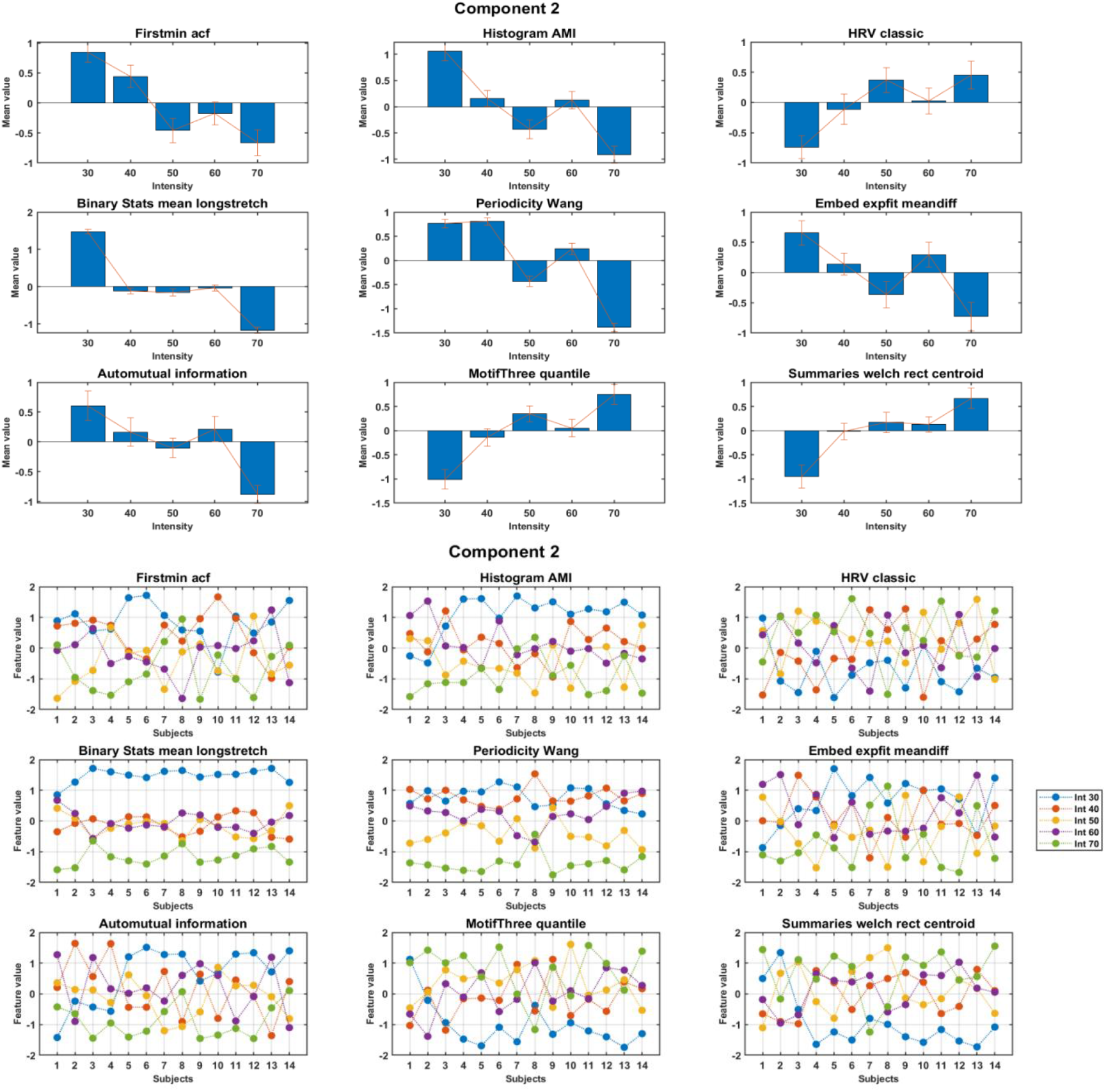
Component 2 with the use of 9 selected features. Upper figure: displaying mean and variance over subjects in each intensity for each feature. Lower figure: displaying the result of the MCCA component for all subjects in each feature.

**Fig S3.**
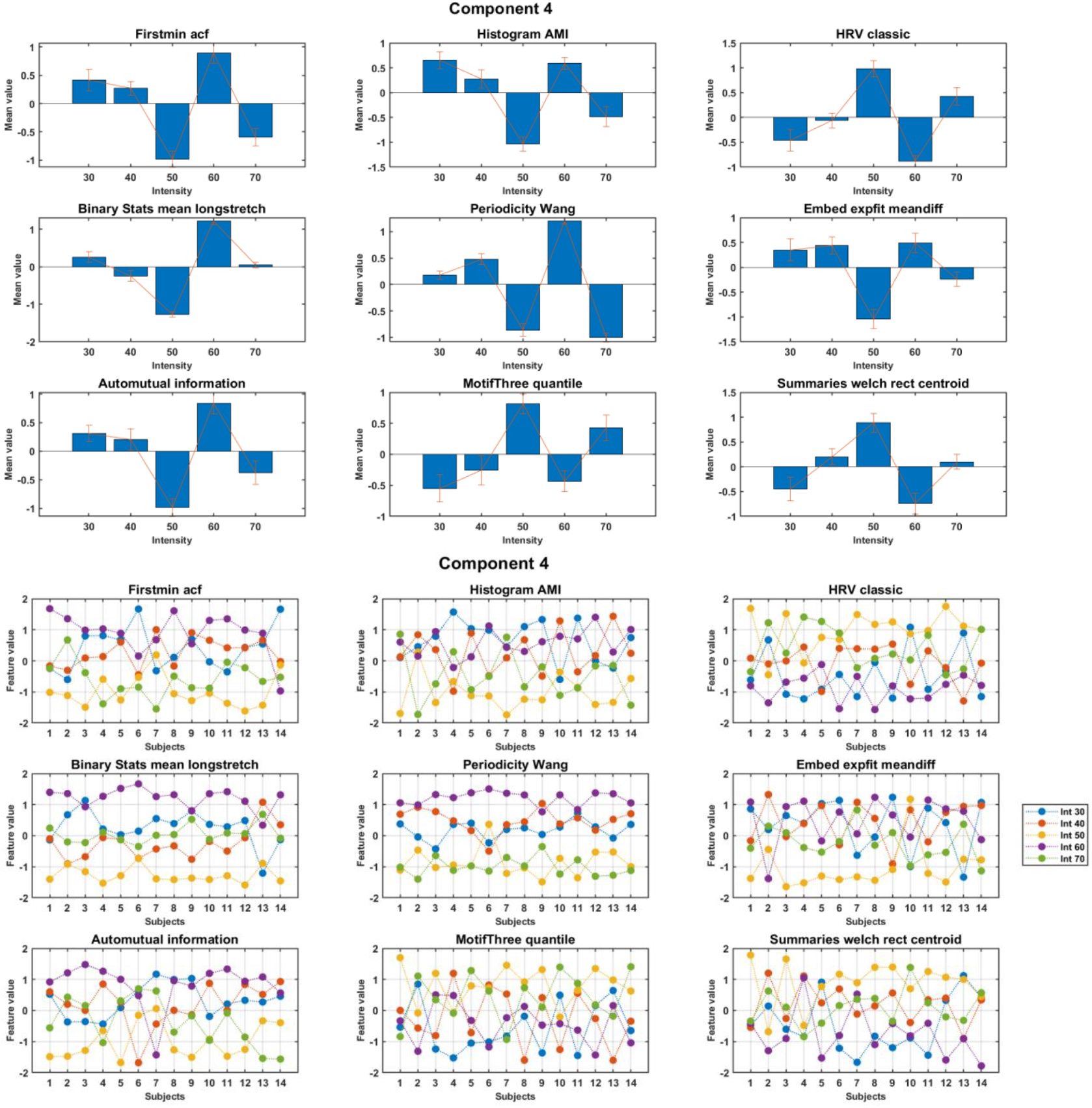
Component 4 with the use of 9 selected features. Upper figure: displaying mean and variance over subjects in each intensity for each feature. Lower figure: displaying the result of the MCCA component for all subjects in each feature.

Supplementary 4: Using the score vector of component 3, we converted the matrix of EEG to a weighted vector. Results are brought in Fig. S4. The shift in dominant frequencies (of weighted averaged signal) from approximately 10 Hz at lower intensities to around 12 Hz at higher intensities demonstrates higher frequency components embedded at common patterns of higher pedaling intensity. This pattern was especially noticeable where the harmonic cluster changed frequency with an increase in intensity. The idea of higher periodicity at elevated intensities is supported by this shift, resulting in a longer time to detect the first peak in the autocorrelation function. This frequency shift suggests that increased physical effort demands more sustained neural processing, potentially reflecting the greater motor coordination required at higher exercise loads.

Supplementary 5: Results of post-hoc analysis for PD-HC comparison and intensity within PD comparison are brought in table S2.

**Fig. S4.**
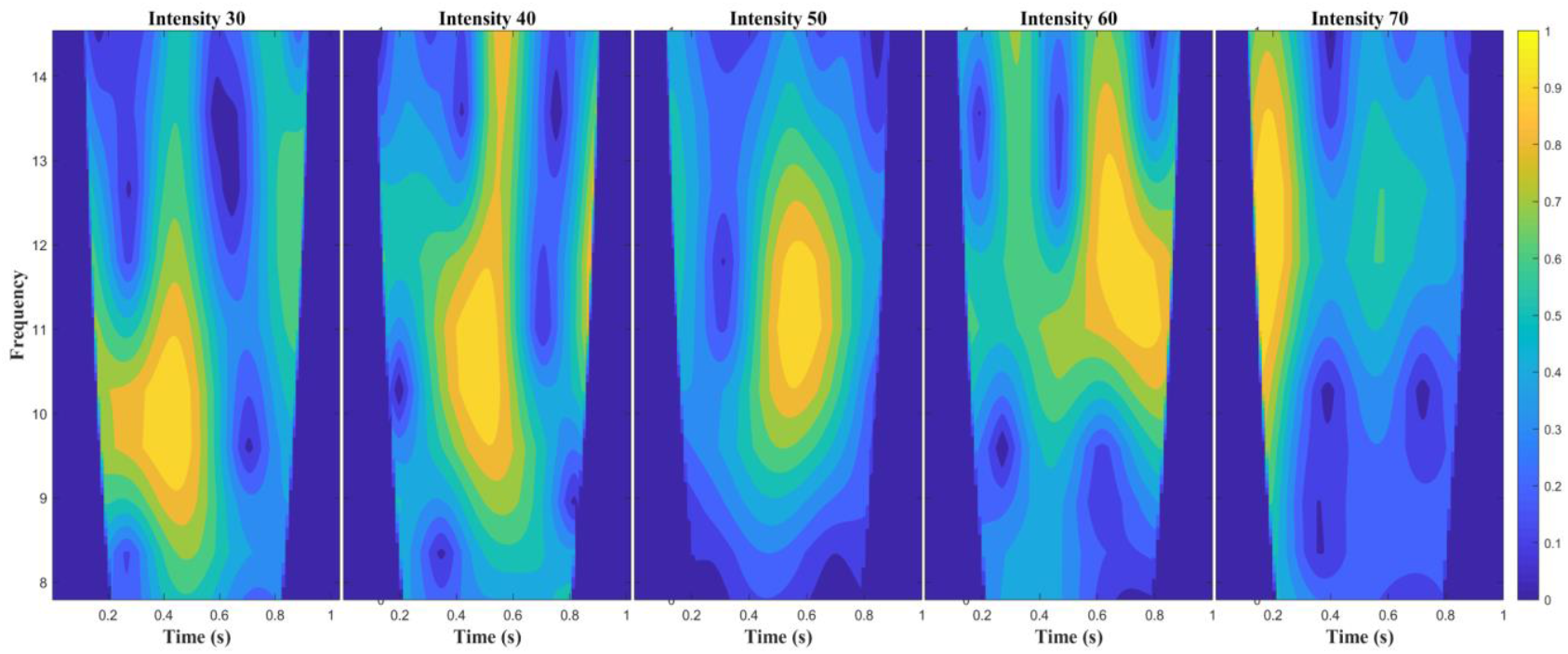
Wavelet of the weighted EEG with MCCA scores for a selected subject, chosen for its clarity and ease of interpretation.

**Table S2:**
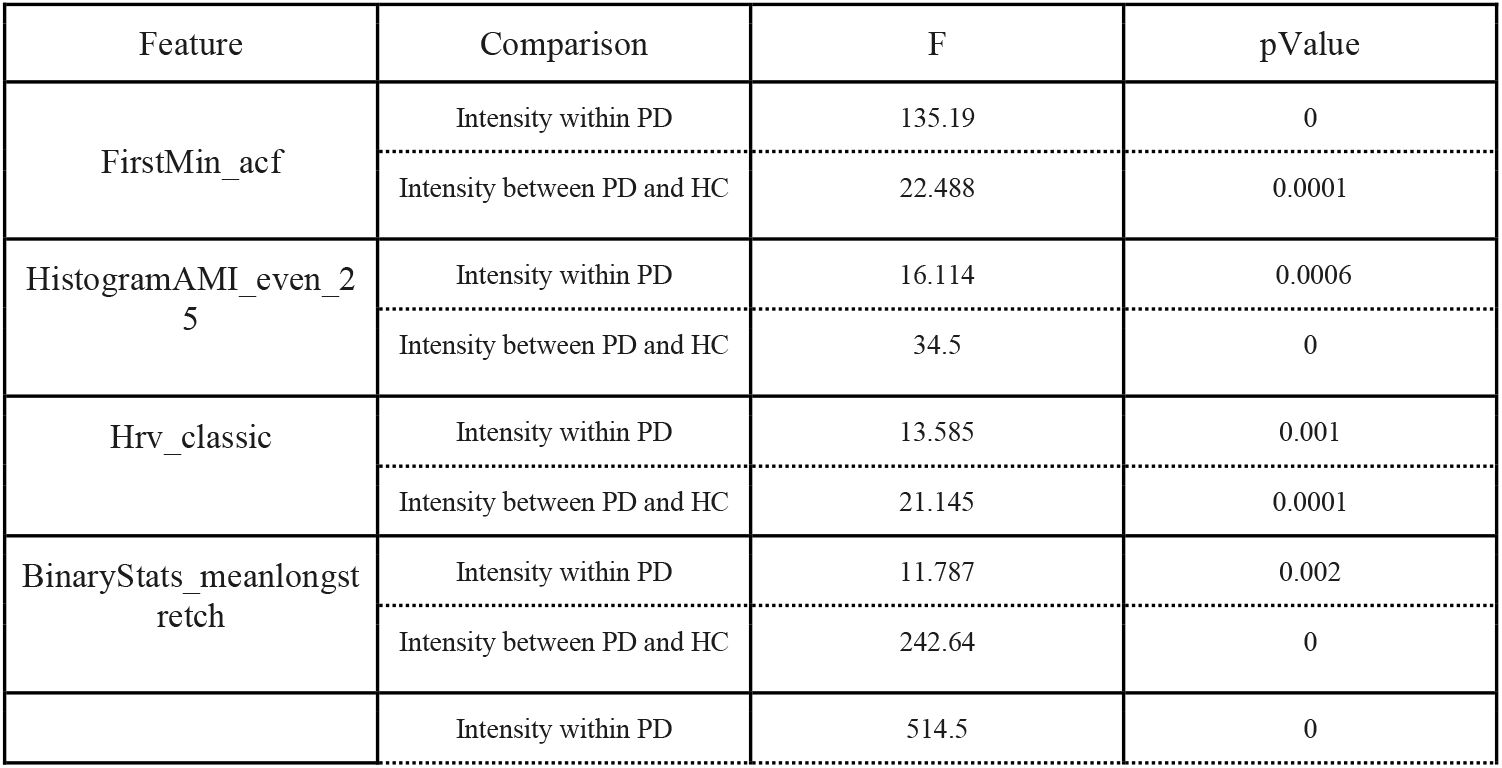
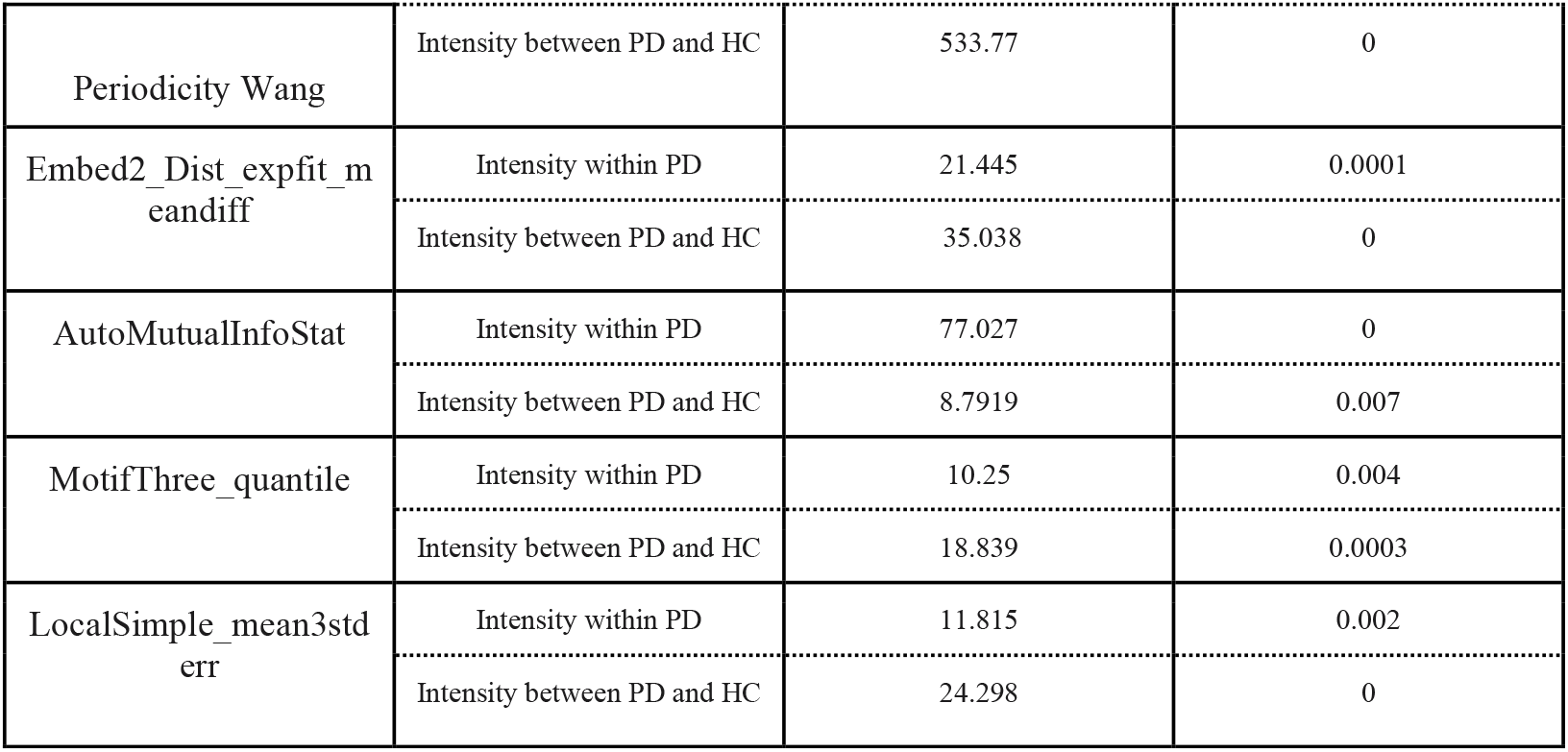
Post-hoc analysis for all 9 important features of component 3.

